# The periphery is the dominant site of B-cell deletion in a polyclonal repertoire

**DOI:** 10.1101/2020.03.10.985192

**Authors:** Jeremy F. Brooks, Raymond J. Steptoe

## Abstract

The concerted actions of multiple tolerance checkpoints limit the possibility of immune attack against self-antigens. For B cells, purging of autoreactivity from the developing repertoire has been almost exclusively studied using B-cell receptor transgenic models. Analyses have generally agreed that central and peripheral tolerance occurs in the form of deletion, receptor editing and anergy. However, when and where these processes occur in a normal polyclonal repertoire devoid of B-cell receptor engineering remain unclear. Here, employing sensitive tools that alleviate the need for B-cell receptor engineering, we track the development of self-reactive B cells and challenge whether deletion plays a meaningful role in B-cell tolerance. We find self-reactive B cells can mature unperturbed by ubiquitous self-antigen expression but, even in the presence of T-cell help, are robustly anergic in the periphery. These studies query the prominence attributed to central and peripheral deletion by most BCR transgenic studies and suggest that other mechanisms predominantly govern B cell tolerance.

## Introduction

Diversity in the primary B-cell repertoire is generated by inherently random V(D)J recombination. While this provides broad protection against pathogens, incidental emergence of autoreactive specificities is common. A series of checkpoints have evolved to minimize the potentially deleterious effects of such B cell receptor (BCR) assembly. These have been defined and extensively explored using B-cell populations with fixed, genetically-modified BCR recognising specific ‘autoantigen(s)’ that have been either randomly knocked-in or targeted to endogenous BCR loci [1]. However, due to technical challenges, repertoires in which self-reactive B cells with diverse clonality and affinities arising in the absence of genetic manipulation of BCR are rarely studied.

Historically, studies have generally agreed that tolerance can be compartmentalised as central and peripheral events, comprising receptor editing, deletion and anergy. Deletion of T and B cells recognising self-antigens at early stages of their development is considered a tenet of self-tolerance [2]. Although deletion has been repeatedly observed in transgenic models of self-reactive B cell development, when and where deletion occurs is unclear across these models. As an example, VH3H9/Vlambda2 B cells reactive with the self-antigen -dsDNA are not eliminated from the repertoire [3], but analogous VH3H9/Vkappa4 B cells with similar specificity are efficiently deleted in the bone marrow [4]. The extent and possibly the site of deletion likely depends on how and where self-antigen is encountered, the affinity of the BCR, and whether there is competition with non self-reactive cells. Recently, studies of polyclonal repertoires where antigen receptors have not been genetically manipulated challenge the primacy of deletion [5-7]. For T cells, the frequency of self-reactive CD8+ve T cells is not different to foreign-reactive CD8+ve T cells and similar numbers of H-Y (male-specific antigen)-binding T cells are present in males and females [5]. These findings are striking given T cells were considered exceptionally sensitive to deletion even when self-antigen was rare [8]. For B cells, while soluble self-antigen can induce deletion in BCR transgenic models, it is insufficient for deletion in a polyclonal repertoire [9]. When immature B cells are surveyed against panels of autoantigens, an unexpectedly large proportion are autoreactive [10, 11] and this persists into the mature B-cell pool.

Here we use tools to directly assay self-reactive B cell development in the absence of BCR manipulation. We report deletion occurs principally in terminally-mature B cells. Residual cells are anergic but do not display those surface markers defined as characteristic of anergy in BCR transgenic models. These findings refine our understanding of integration of into the development and control of self-reactive B cells.

## Methods

### Mice

Actin.mOVA (act.OVA) mice ubiquitously express membrane-bound ovalbumin (OVA) under the control of the β-actin (CAG) promoter [12]. Act.OVA mice were maintained as heterozygous x null matings with C57BL/6LArc mice sourced from Animal Resources Centre (Perth, WA Australia). UBC.mHEL3x mice ubiquitously expressing membrane-bound hen egg lysozyme [13] under the control of the human ubiquitin C promoter and control C57BL/6JAusb were a gift from Professor Robert Brink. C57BL/6JArc mice were purchased from the Animal Resources Centre. All mice were bred and/or maintained at the TRI Biological Resources Facility (Brisbane, Australia). Male and female, age-matched (typically within 4-6 weeks) transgenic and non-transgenic littermates 6-16 weeks of age were used unless stated otherwise. Studies were approved by the University of Queensland Ethics Committee.

### Flow cytometry

Pooled lymph nodes (inguinal, brachial, axillary, mesenteric) and spleens for naïve mice or inguinal lymph nodes from immunized mice were collected and pressed through a 70μM strainer (Corning, NY, USA) to create a single cell suspension. For bone marrow, femurs and tibiae were harvested and BM flushed with cold PBS/FCS (2.5% v/v). BM was then passaged through a 21g needle to prepare a single cell suspension. Erythrocytes were lysed (spleen and BM) using ammonium chloride buffer. Antibody and tetramer staining were performed on ice as described [14]. Absolute cell number was enumerated using a bead-based counting assay with Flow-Count™ fluorospheres (Beckman Coulter, CA, USA) as described [14]. IgM-PECy7 (RMM-1), IgD-BUV395 (11-26c.2a), CD4-FITC (GK1.5), CD4-PECy7 (RM4-5), CD4-PE (RM4-4), CXCR5-APC (J252D4), CD8-PerCPCy5.5 (53-6.7), CD8-FITC (53-6.7), CD8-PECy7 (53-6.7), CD11c-FITC (N418), CD11c-PE (N418), B220-APC (RA3-B62), CD19-APC (6D5), CD19-BV785 (6D5), CD21/35-Pacific Blue (7E9), PD-1-BV785 (29F.1A12), CD23-AF700 (B3B4), B220-FITC (RA3-6B2) and GL7-Pacific Blue (GL7) were from Biolegend (CA, USA). CD93-BV650 (R139), were from BD Bioscience (CA, USA). NK1.1 (PK136), Gr1 (RB6-8C5) and CD11b (M1-70) were purchased from BioXCell (NH, USA) or grown in-house and labelled with FITC in house. For tetramers, OVA and HEL were biotinylated in house to ∼1 biotin per antigen molecule and tetramers constructed as described [14] using streptavidin-PE (Biolegend, CA, USA), streptavidin-APC (Biolegend, CA, USA), or streptavidin-BV510 (BD Bioscience, CA, USA). Data were collected on an LSR Fortessa X-20 (BD Bioscience, CA, USA) and analysed using Diva Version 8 (BD Bioscience, CA, USA) or FlowJo Version 10 (Flowjo LLC, CA, USA).

### Immunizations and ELISA

OVA (Grade V, Sigma, MO, USA) or PBS was emulsified in incomplete Freund’s adjuvant (Sigma MO, USA) using a mini-BeadBeater (Biospec Products OK, USA) and 100μL injected subcutaneously at the tail base. For SA-OVA, biotinylated OVA was mixed in a 1:1 ratio with SA (Promega, WI, USA) and emulsions were prepared as described. OVA- and SA-specific IgG1 ELISA was as described [14] using serial 4-fold dilutions of plasma.

### Statistical analysis

Data was tested for normality and means compared using student t-test with multiple testing correction where required or ANOVA with Tukey post-hoc analysis for normally-distributed data and Mann-Whitney test or Kruskal-Wallis test for non-normally distributed data Antibody titres were log-transformed for statistical comparisons. (GraphPad Prism V8.01, GraphPad Software, Inc. – CA, USA).

## Results

### Self-reactive B cells are unexpectedly common in the peripheral B cell pool

We used a multi-tetramer approach understand B-cell selection events in the periphery where antigen-specific B cells were present at a natural frequency without genetic modification of the BCR. We assessed development of ovalbumin (OVA)-specific B cells in a mouse model where membrane-bound OVA is ubiquitously expressed (act.OVA) **(Suppl 1 A,B**) and compared development in OVA-ve (non-Tg) littermates.

We first probed spleen as the initial peripheral site of terminal B-cell maturation. In line with established interpretations [15], we considered that numerical differences in antigen-specific B cells within the transitional population would reflect central selection events in BM. Among IgM/IgD-defined transitional B cells, the number of OVA-specific cells did not differ between OVA+ve mice and OVA-ve littermate controls **(Fig. 1A,B)**, indicating that clonal deletion had not occurred in the BM nor, prior to this, in the periphery. Similarly, the numbers and frequency of OVA-specific mature B cells (**Fig. 1C**) did not differ. In ascertaining OVA-specific B-cell frequency in lymph nodes (LN) **(Fig.1D,E)** as a site of B-cell traffic following terminal development, we did, however, find significantly fewer OVA-specific B cells in OVA+ve mice than in OVA-ve littermate controls **(Fig. 1D,E)**. In keeping with otherwise normal B cell development in these mice, the overall populations of splenic transitional (IgM^+^IgD^lo^) and mature (IgM^var^IgD^+^) B cells and mature LN B cells were similar in number and proportion between OVA+ve and littermate control mice **(Suppl 1C-G)**.

**Figure 1.**
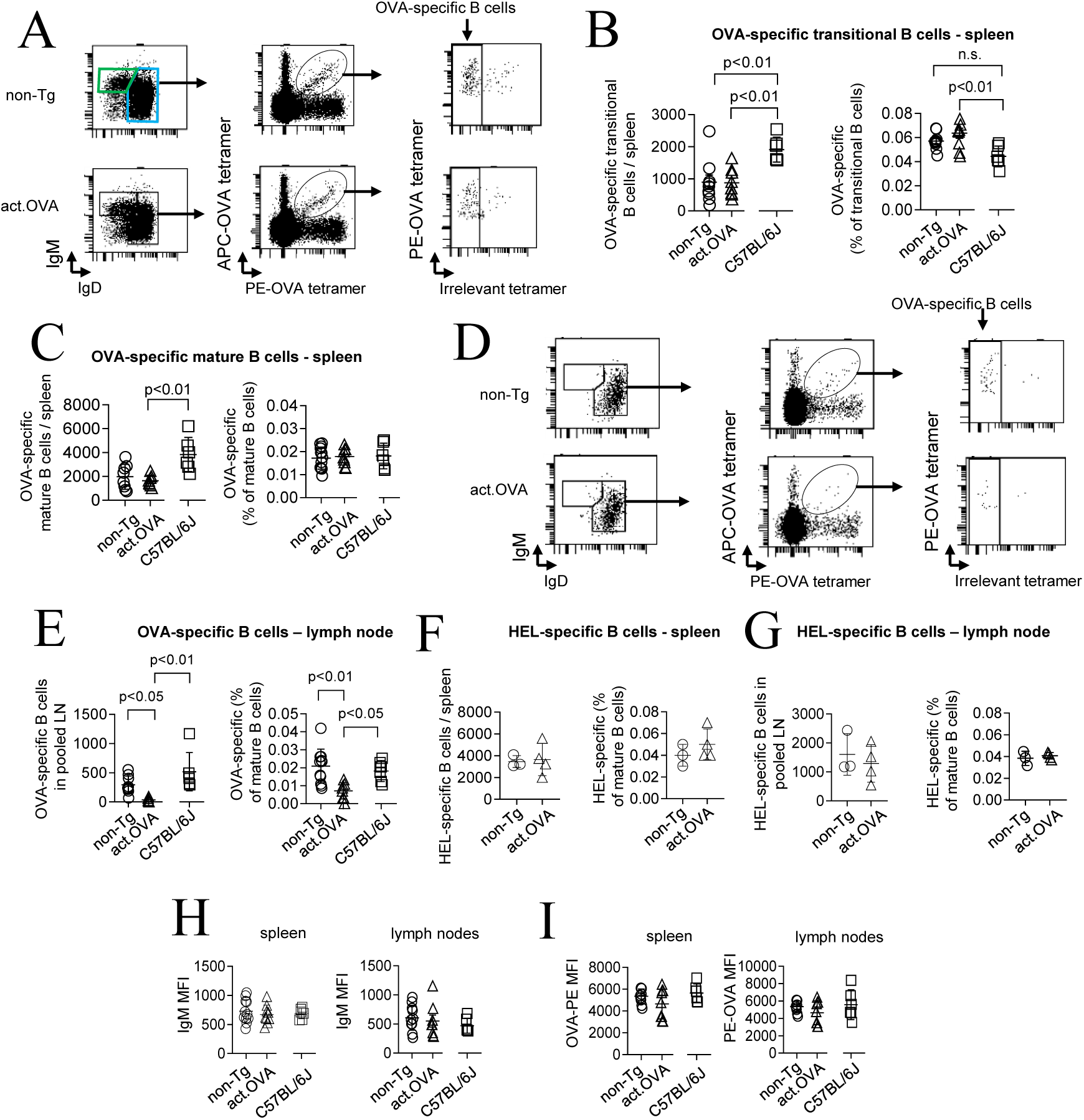
Unexpected retention of self-reactive B cells in the periphery. Spleens or LN (inguinal, axillary, brachial, mesenteric) were harvested from non-Tg, act.OVA or C57BL/6 mice and stained with OVA or HEL tetramers. **A**) Representative analysis of mature (blue gate) or transitional (green gate) B cells with OVA tetramer binding. **B,C**) OVA-specific B cell number and frequency among splenic transitional (**B**) and mature (**C**) B cells. **D**) Representative analysis of LN B cells. **E**) OVA-specific B cell number and frequency in pooled LN. **F,G**) HEL-specific B cell number and proportion among splenic (**F**) and pooled LN (**G**) mature B cells (**F**). **H, I**) IgM expression (**H**) and OVA-PE tetramer staining intensity (**I**) on mature B cells in spleen and LN. Data show representative plots or individual mice (mean±SD) pooled from 2-5 experiments. ANOVA with Dunnett’s post-test (B,C,E,), Kruskal-Wallis (E, left panel) or t-test (F,H,I).

In selecting OVA-ve control mice we also analysed OVA-specific B-cell frequency in the C57BL/6JArc mice used to maintain the act.OVA colony. Interestingly, we observed more OVA-specific B cells in both spleen and LN of C57BL/6JArc controls than in OVA-ve littermates **(Fig. 1C,E)**, but this was an artefact due to a larger B cell compartment in externally-sourced C57BL/6JArc mice **(Suppl Fig. 1H**). Within the total B cell pool OVA-specific B-cell frequency did not differ between C57BL/6JArc and OVA-ve littermates **(Fig. 1C,E; Suppl 1I)** confirming this and emphasising the necessity of selecting appropriate controls. Using C57BL/6 from an alternative housing source rather than mice from the same facility could lead to the erroneous conclusion that OVA-specific deletion had occurred. This may account for the reports of apparent deletion in a previous study of act.OVA mice in which act.OVA and non-Tg C57BL/6J controls were sourced from different locations [9]. The total number and frequency of hen-egg lysozyme (HEL)-specific B cells in spleen and LN, as an alternative (non self-reactive) specificity, did not differ between OVA+ve and OVA-ve littermates (**Fig 1F,G)**, allowing us to conclude the reduced frequency of OVA-specific B cells in LN of OVA+ve mice was a consequence of peripheral OVA expression rather than a non-specific effect.

Down-regulation of surface IgM on self-reactive cells that escape deletion is reported as a canonical feature of tolerance in many BCR-Tg repertoires [1]. However, in agreement with Taylor et al. [9], we found no difference in expression of surface IgM or IgD on OVA-specific B cells between OVA+ve and OVA-ve littermate controls in either spleen or LN **(Fig. 1H; Suppl 1J)** and they bound equivalent amounts of tetramer **(Fig. 1I)**. This is consistent with previous assessments of polyclonal repertoires and some monoclonal BCR-Tg systems that propose deletion is not the dominant mechanism of central tolerance [9, 11, 16]. Therefore, deletional tolerance appeared to be operating in the periphery, but at a much later developmental stage than previously appreciated.

### Compartmentalising naïve B cell subsets confirms deletion is restricted to the lymph nodes

While a broad analysis indicated peripheral deletion manifested only among mature LN B cells, it is possible deletion occurred within a sub-population of splenic B cells and was not detected by a broad analysis. When we more deeply probed splenic B-cells subpopulations (**Fig. 2A**), none harboured lower numbers of OVA-specific cells in OVA+ve mice **(Fig. 2B**), nor did the proportions differ (**Suppl 2A).** In contrast, within the mature follicular B-cell population in LN, OVA-specific B cells were significantly reduced in OVA+ve mice in both number **(Fig. 2B)** and proportion (**Suppl Fig. 2A**), but this did not manifest as a change when all B cell subpolulations were pooled (Fig. 2B),confirming the conclusions of the broad analysis.

**Figure 2.**
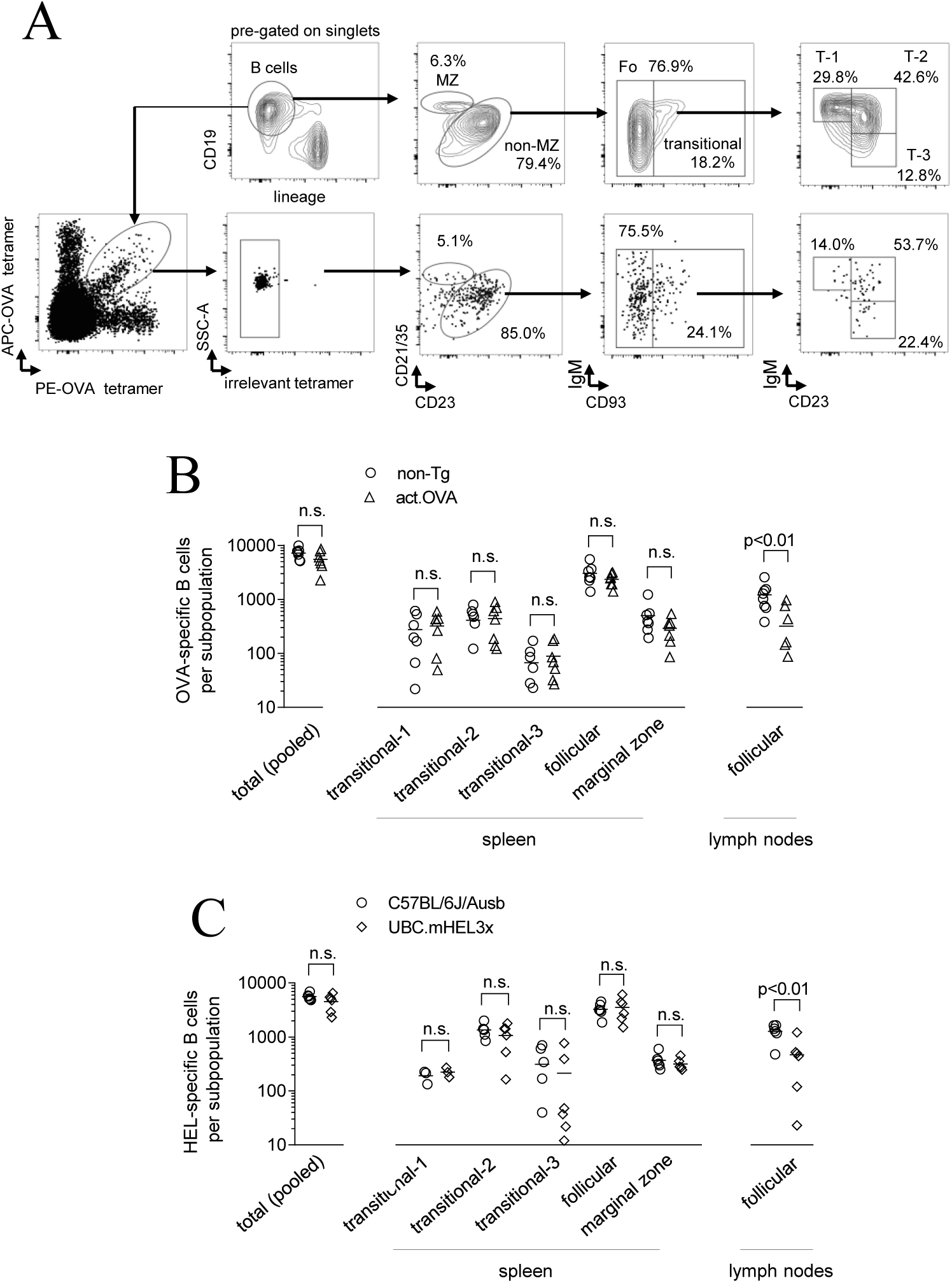
Lymph nodes are a major site for deletion of self-reactive B cells. Spleen and lymph nodes were harvested from act.OVA mice and non-Tg littermates, or UBC.mHEL3x and C57BL/6JAusb mice, and stained for OVA-specific and HEL-specific B cells respectively. A) Gating strategy to identify peripheral B cell subsets. B) OVA-specific B cell numbers for subpopulations in spleen and follicular B cells in lymph nodes. Total (pooled) is sum of all OVA-specific B cells detected. C) As in B, for HEL-specific B cells. Data show representative FACS plots from (A) or individual mice (B,C) pooled from 2 separate experiments (mean±SD). T-test with multiple testing correction.

To determine whether peripheral deletion at a late stage of maturation was a universal pattern we studied UBC.mHEL3x mice which ubiquitously express the widely studied antigen HEL as an alternative to OVA. Detailed analysis (**Fig. 2A**) showed no difference between HEL+ve mice and HEL-ve controls in either the number (**Fig. 2C**) or proportion (**Suppl 2B**) of HEL-specific B cells within subpopulations of splenic B cells. Importantly, we found the number (**Fig. 2C**) and proportion (**Suppl 2B**) of HEL-binding follicular B cells was, however, significantly reduced in LN of HEL+ve mice despite similar numbers of B cells within each compartment **(Suppl 2D)** showing that UBC.mHEL3x mice phenocopied act.OVA mice. Here, because we used native HEL to detect HEL-specific B cells in HEL3x mice, which express HEL with 3 point mutations that affect binding by the HyHEL10 mAb but where overall protein folding and binding to other anti-HEL mAb is not affected [17], we may have underestimated slightly the extent of deletion. Nevertheless, our data confirm that self-reactive B cells in a repertoire devoid of an engineered BCR remain mostly unperturbed by deletion until late in their development.

### A shadow of peripheral B cell deletion is present in bone marrow, but only in recirculating mature B cells

Our analyses thus far revealed self-reactive B-cell deletion is restricted to terminally-mature cells, possibly after transiting spleen, and central deletion in BM is less apparent than in BCR-Tg models. We pondered whether directly assessing BM B cells might reveal centrally-mediated deletion. In BM, IgM and IgD readily separated developing IgM^hi^IgD^lo^ immature B cells from recirculating mature B cells **(Fig. 3A)**. Tetramers were sufficiently sensitive to reveal small populations of OVA-binding (**Fig. 3A**) or HEL-binding B cells, but not in those devoid of surface Ig (i.e. IgM^-^IgD^-^) **(Fig. 3A)**. Interestingly, among recirculating mature B cells, the number of OVA-binding and HEL-binding cells was markedly reduced in act.OVA and UBC.mHEL3x mice, respectively, relative to controls **(Fig. 3B,C)**. By contrast, no such differences were present among immature IgM^hi^IgD^lo^ B cells **(Fig 3D,E)**. These data reinforce the finding that antigen-specific deletion manifests only among terminally-mature B cells in the periphery.

**Figure 3.**
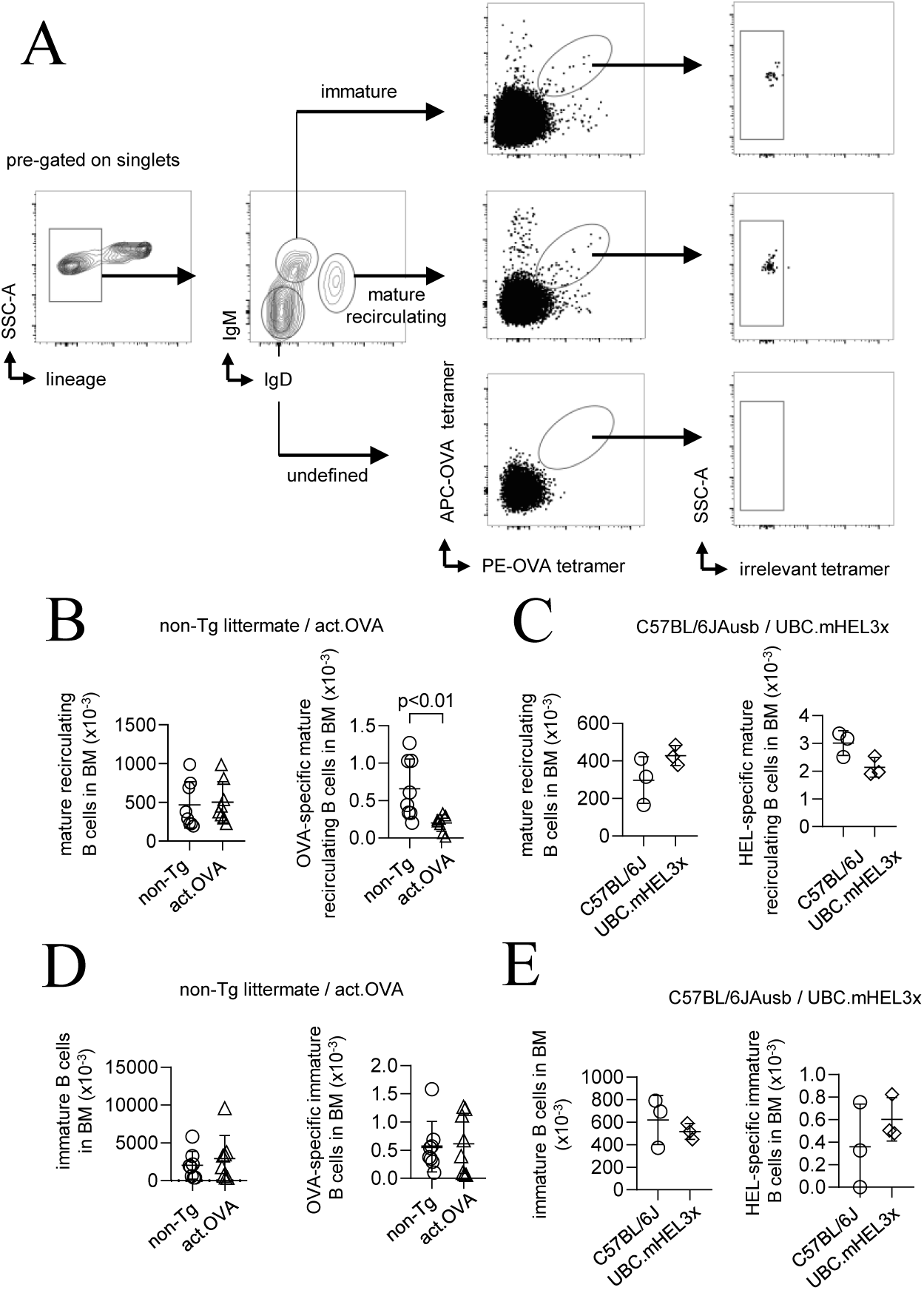
Self-reactive B cells are deleted from the recirculating pool of mature B cells. Bone marrow was harvested from act.OVA and non-Tg littermates or UBC.mHEL3x and C57BL6JAusb mice and probed with OVA or HEL tetramers, respectively. A) Gating strategy to identify OVA-specific cells among immature and mature B cells in bone marrow. B) Number of mature B cells (left) and OVA-specific mature B cells (right). C) Number of mature B cells (left) and HEL-specific mature B cells (right). D) Number of immature B cells (left) and OVA-specific immature B cells (right). E) Number of immature B cells (left) and HEL-specific immature B cells (right). Data are representative FACS plots (A) or individual mice (mean±SD) (B,C,D,E), pooled from 2 experiments (except C,E which was a single experiment, n=3) t-test(B,D,E).

### Age does not substantially impact the peripheral self-reactive pool

Age-related changes in the B-cell compartment appear to enrich for autoreactivity and humoral dysfunction as life progresses [18, 19]. Because the repertoire of young mice expressing either OVA or HEL harboured a significant population of self-reactive cells, we predicted this may increase with age. We probed the spleen and lymph nodes of mice aged for at least 8 months. In spleen, although we only assayed a small number of act.OVA and UBC.mHEL3x mice, we found that the frequency of OVA- or HEL-specific B cells had not substantially changed since youth **(Fig 4A,B, Supplementary Fig 3F, G)**, where the frequency remained relatively equal between non-Tg and act.OVA mice **(Fig 3A)**. In contrast, in the lymph nodes, we no longer detected deletion of OVA-specific B cells in act.OVA mice **(Fig 3A)**. We observed a similar trend in the spleen and lymph nodes of UBC.mHEL3x mice **(Fig 3B)**. The data, from a limited number of mice, suggest that self-reactive B cells persist in the peripheral pool and are continuously exported from the bone marrow throughout life.

**Figure 4.**
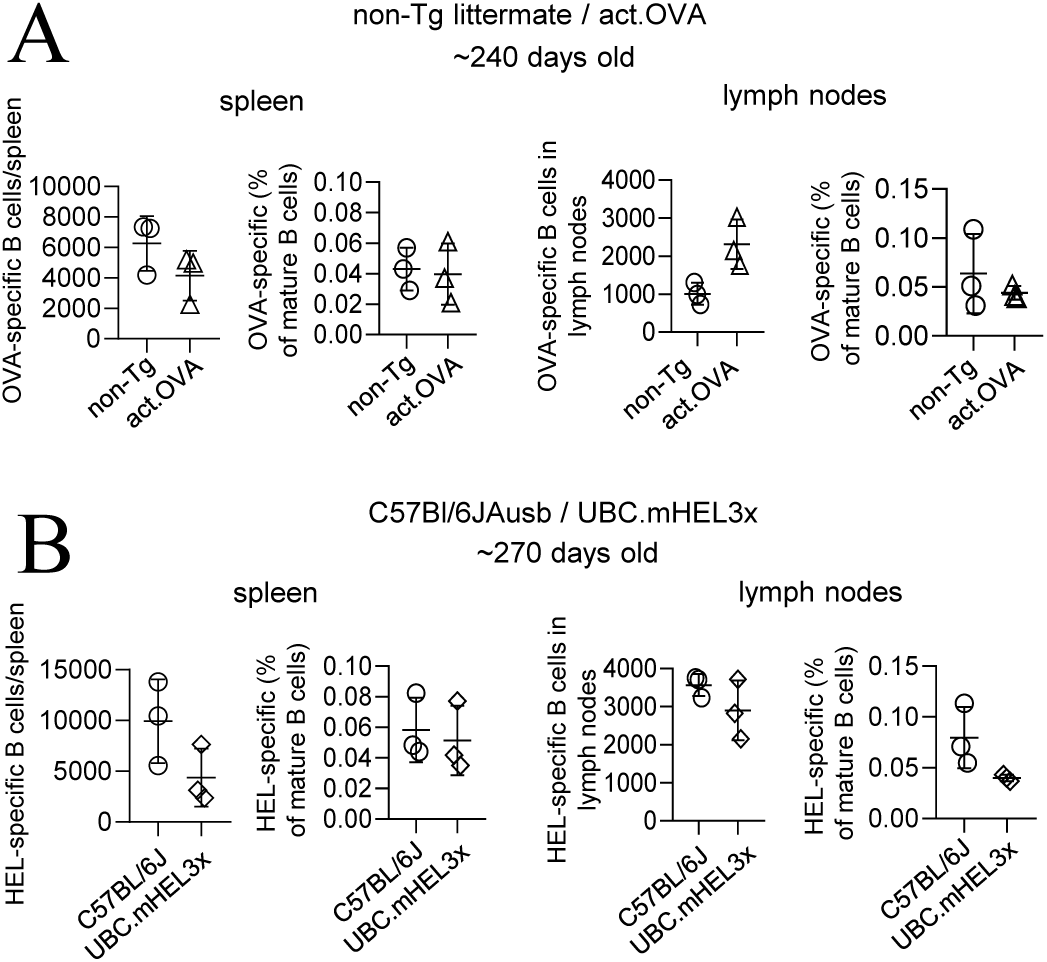
Self-reactive B cells persist in the periphery throughout life. Spleen and LN (inguinal, axillary, brachial, mesenteric) were harvested from act.OVA mice and non-Tg littermates, or UBC.mHEL3x and C57BL/6JArc mice, at the ages shown and stained for OVA-specific and HEL-specific B cells respectively. A) Number or proportion of OVA-specific B cells in spleen and LN. B) Number or proportion of HEL-specific B cells spleen and LN. Data show individual mice pooled from two experiments (mean±SD).

### Self-reactive cells that escape deletion are controlled principally by anergy

The limited deletion of self-reactive B cells we defined in a normal polyclonal repertoire emphasizes the importance of peripheral non-deletional B-cell checkpoints to restrain autoantibody production. Immunization of act.OVA mice with OVA/IFA led to little OVA-specific IgG1 production compared to non-Tg littermates (**Suppl 3A**) suggesting OVA-specific B cells were anergic or kept in check by other cell-intrinsic mechanisms. As this might reflecting T-cell tolerance present in act.OVA mice [12], we designed experiments to bypass this **(Fig. 5A)**. As anergic self-reactive B cells present antigen to T cells as efficiently as non-anergic cells [20] we immunized mice with a conjugate of streptavidin and OVA (SA-OVA) in IFA. Here, OVA-specific B cells take up SA-OVA and present both SA- and OVA-derived peptides enabling T-cell help from SA-specific T cells in act.OVA mice **(Fig. 5A)** without the need for adoptive T cell transfer or red cell carriers as used in other settings [9, 21].

**Figure 5.**
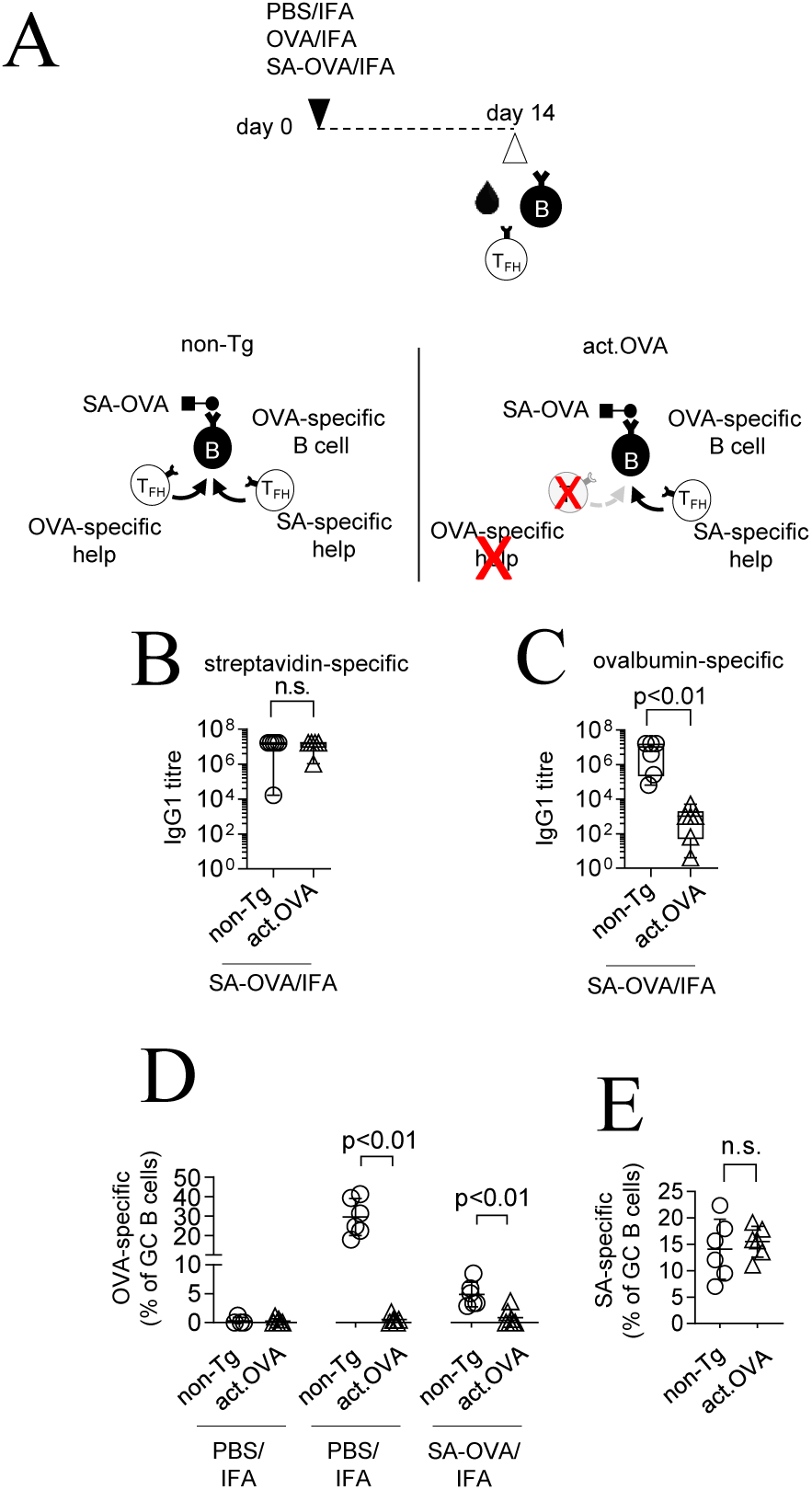
Residual self-reactive B cells are functionally anergic. A) Experimental layout. B-E) Non-Tg and act.OVA mice were immunized with OVA/IFA, SA-OVA/IFA or PBS/IFA and two weeks later plasma was assayed for IgG1 (B) or draining (inguinal) LN tetramer analysed by flow cytometry for OVA-Specific GC B cells (C) or SA-specific B cells (D). Data show values for individual mice (mean±SD) pooled from two (B) or 4 (C,D) independent experiments. Mann-Whitney test of log-transformed titres (B,C) Mann-Whitney (D) or t-test (E).

Few CD4+CXCR5+PD-1+ Tfh were present in sham-immunised mice, **(Suppl 3B,C)** and this was similar in OVA-immunised act.OVA mice consistent with OVA-specific T tolerance **(Suppl 3B,C)**. However, Tfh were strongly and equivalently elicited by SA-OVA/IFA in act.OVA and non-Tg littermates (**Suppl 3B,C**). *In vitro* restimulation assays showed that OVA/IFA and SA-OVA/IFA had induced cytokine production in OVA-responsive T cells only in non-Tg littermates whereas OVA-SA induced SA-responsive cytokine producing T cells in both act.OVA and non-TG littermates, validating the immunisation approach (**Suppl 3D**). SA-OVA/IFA immunization elicited similar high titres of anti-SA IgG1 in OVA+ve mice and non-Tg littermates **(Fig. 5B)** further indicating the effectiveness of SA-specific T-cell help Conversely, SA-OVA/IFA elicited only limited amounts of anti-OVA IgG1 in act.OVA mice (**Fig. 5C**) demonstrating that provision of linked SA-specific help did not restore OVA-specific IgG1 productionthereby indicating that anergy or other cell-intrinsic mechanisms substantially limited OVA-specific B-cell responsiveness. Consistent with this, the frequency and number of OVA-specific GC B cells that developed after OVA/IFA immunization was limited in act.OVA mice (**Fig. 5D, Suppl 3F,G**) and OVA-specific GC B cell differentiation in act.OVA mice was not restored by provision of help through SA-OVA immunization (**Fig. 5D, Suppl 3H**) even though SA-specific GC B cell development was substantial regardless of the presence of OVA (**Fig. 5E, Suppl 3I**).

## Discussion

Because some important biological functions rely on the retention of autoreactive B cells, the findings described here may not be entirely surprising. For example, pathogens that express self-like epitopes cannot be resisted by the host if the repertoire is devoid of B cells recognising self-antigen. That B cells can re-diversify their BCR inside the GC led to the ‘clonal redemption’ hypothesis that proposes rare self-reactive B cells can be counterselected for binding to self-antigen while maturing affinity to cross-reactive self-like epitopes that are expressed by pathogens [22]. Clonal redemption has now been formally demonstrated in both mouse [21] and man [23] underscoring a need to preserve at least some degree of autoreactivity in the B cell repertoire. Our study shows that these self-reactive B cells are more common than previously thought.

In a landmark study that characterised self-reactivity among polyclonal B cells, Wardemann and colleagues showed for the first time how polyclonal self-reactive cells develop in the repertoire [11]. Their approach screened monoclonal antibodies generated from B cells at different developmental stages in the repertoire against a defined panel of autoantigens. Of the cells cloned, many of them had arrived in the periphery with self-reactivity and could mature without being deleted. A more high-throughput assay recently confirmed these observations [10]. The findings from those methods are supported by the Nur77-GFP mouse model that faithfully reports BCR signaling [24]. Of mature cells in the periphery, many were GFP+ at steady state, presumably in response to endogenous self-antigen.

To our knowledge, only one other study has directly assessed deletion of polyclonal autoreactive B cells as opposed to indirectly (cloning, reporter assays). Taylor et al used an approach similar to the one described here [9]. OVA-specific B cells which bound tetramers were enriched and characterised. When OVA was expressed as a soluble self-antigen, no deletion was detected. When OVA was expressed as a membrane-bound self-antigen, modest deletion (about half of the total number) was detected in the periphery. The discrepancies between our findings are likely methodological, as we have used a direct staining approach that does not require enrichment and, where possible, have accompanied our analysis with the most appropriate negative controls. Independent studies using similar enrichment methods show insulin-binding B cells, a bona fide autoreactive specificity, are also common among mature B cells [25]. In subsequent studies, the inhibitory signalling pathways underpinning anergy in autoreactive cells are actually elevated among much of the bulk mature follicular B cell compartment, leading the same laboratory to conclude that autoreactivity is more common in the repertoire than previously appreciated [26].

Recent studies on deletional tolerance in T cells report analogous discrepancies between TCR transgenics and polyclonal T cells [5-7]. It was previously thought that T cells were exceptionally sensitive to deletion even at low levels of self-antigen expression [8]. However, enumerating polyclonal T cells specific for H-Y, which were predicted to be deleted from male mice that express thymic Hy antigen [27], showed the frequency is not different to female littermates [5]. In agreement, enumerating GFP-specific T cells in mice expressing GFP as a self-antigen under various tissue-specific promoters revealed that deletion can be minimal even when self-antigen is widely expressed [6]. Together with our data here on B cells, we argue that substantial deletion observed in engineered immune repertoires may largely be explained by nonphysiological frequency and affinity of autoreactive T and B cells.

An important question that remains is how receptor editing, which appears to dominate over deletion during central selection [28], integrates tolerance in the emerging repertoire. Further studies should aim to sort and clone individual autoreactive B cells from a defined population, as shown here, to determine the degree of receptor editing. It is possible that although similar frequencies of autoreactive cells develop in act.OVA mice, these cells have lowered affinity for monomeric OVA endowed by receptor editing and this helps to maintain anergy.

How easily can anergy be broken to enable pathogenesis (against self) or protection (against self-like foreign)? We and others have shown that self-reactive B cells can become transiently activated, even on a non-autoimmune murine background, when highly immunogenic stimuli are supplied at the same time as self-antigen encounter (manuscript submitted). This setting may reproduce what happens during inflammation, where self-reactive B cells solicit not only local T cell help in trans but also diverse innate signals that bolster cells past an activation threshold required for autoantibody production. While transient activation probably occurs frequently in the developed repertoire, extrinsic immune regulation (e.g. regulatory T cells) and additional peripheral checkpoints (e.g. Fas) also limit the effector function of autoreactive B cells. Mutations in those additional tolerance checkpoints would permit less stringent control of these anergic B cells [22, 29]. Evidence of this can be drawn from neutralising antibodies against HIV which are isolated from hosts with ‘relaxed’ B cell tolerance and often originate from germline autoreactive precursors [30].

In conclusion, we provide a new analysis for a question which still remains to be resolved – how are autoreactive B cells kept in check? Deletion does not appear to operate as a tolerance mechanism for B cells until late in their development and only a small fraction goes on to be removed. Given this discrepancy with BCR transgenic mice, our data underscore the need to explore tolerance mechanisms active in the endogenous repertoire without genetic engineering of the BCR.

## Acknowledgments

The authors thank Professor Robert Brink for providing UBC.mHEL3x mice, Professor Marc Jenkins for providing act.OVA mice, and the TRI flow cytometry core for outstanding technical support.

---

**Supplementary Figure 1.**
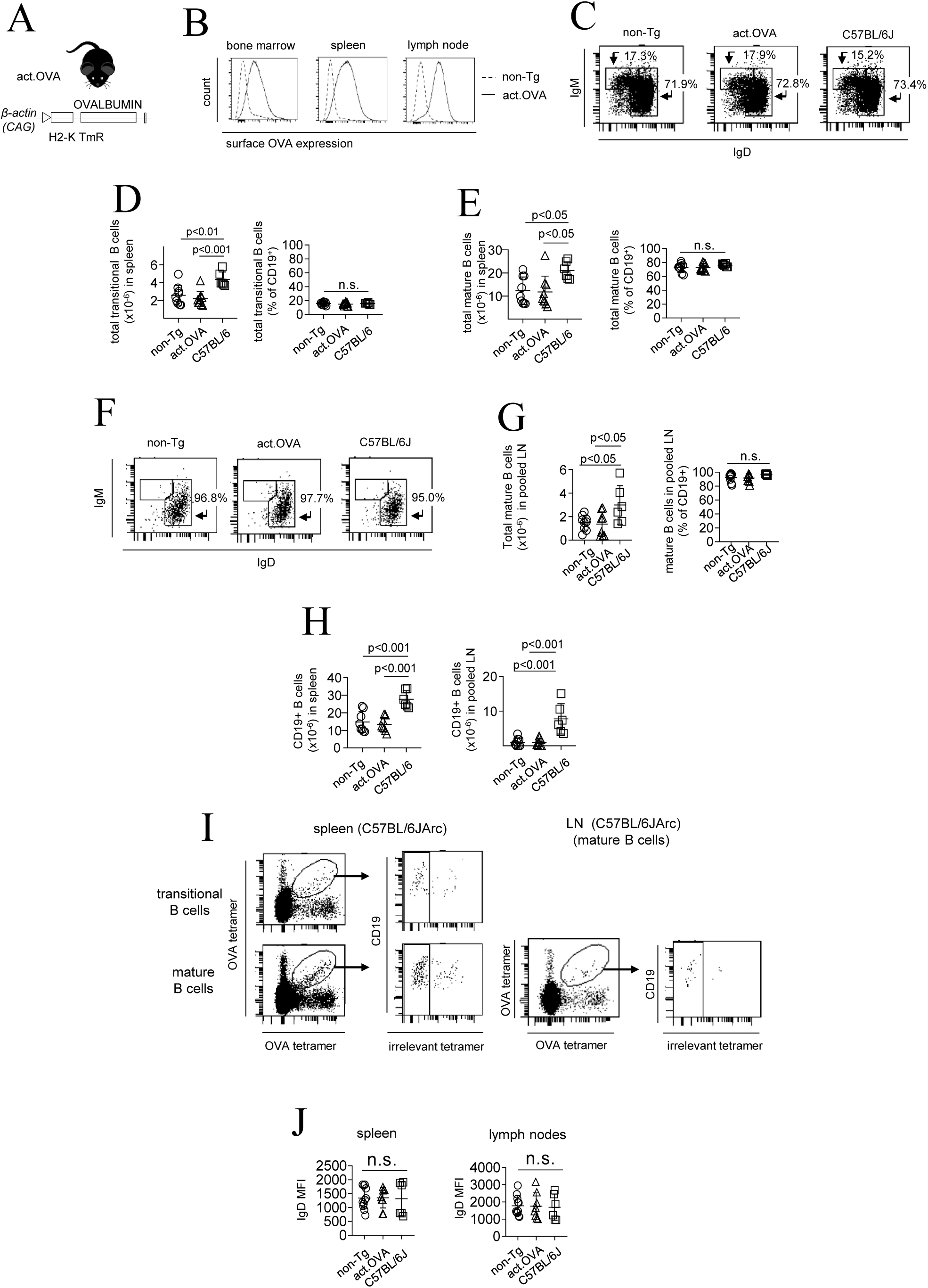
Development of OVA-specific B cells in act.OVA mice. Spleen or pooled (inguinal, brachial, axillary, mesenteric) LN were harvested for analysis from act.OVA (OVA+) mice and non-Tg (littermate) controls. In some cases C57BL/6JArc mice were also analysed. A) Schematic of act.OVA mouse. B) Staining for surface OVA expression in bone marrow (central compartment), spleen and lymph nodes (peripheral compartments). Single cell suspensions were stained with an anti-OVA rabbit polyclonal antibody (Sigma) followed by a biotinylated polyclonal Donkey anti-rabbit secondary antibody (Merck). Cells were then stained with SA-FITC and analysed by flow cytometry. C) Representative flow plots showing IgM/IgD expression and gating for transitional and mature B cells in spleen. Inset values are the proportion of gated populations as a percentage of CD19+ B cells. D) Total number (left) and percentage (right) of transitional B cells in spleen. E) Total number (left) and percentage (right) of mature B cells in spleen. F) Representative flow plots showing IgM/IgD expression and gating of B cells in LN. Inset values are the proportion of the gated populations as a percentage of CD19+ B cells. G) Total number (left) and percentage (right) of mature B cells in LN. H) Total number of B cells in spleen (left) and LN (right) of act.OVA, non-Tg (littermate) and C57BL/6JArc mice. I) Representative staining for OVA-specific B cells in transitional and/or mature B cell compartments within the spleen (left) and LN (right) of C57Bl/6JArc mice. J) Surface IgD staining OVA-specific B cells in spleen and LN. Data are representative dot plots (C,F,I) or individual values (mean±SD) pooled from 4-5 experiments. ANOVA with Tukey’s post-test.

**Supplementary Figure 2.**
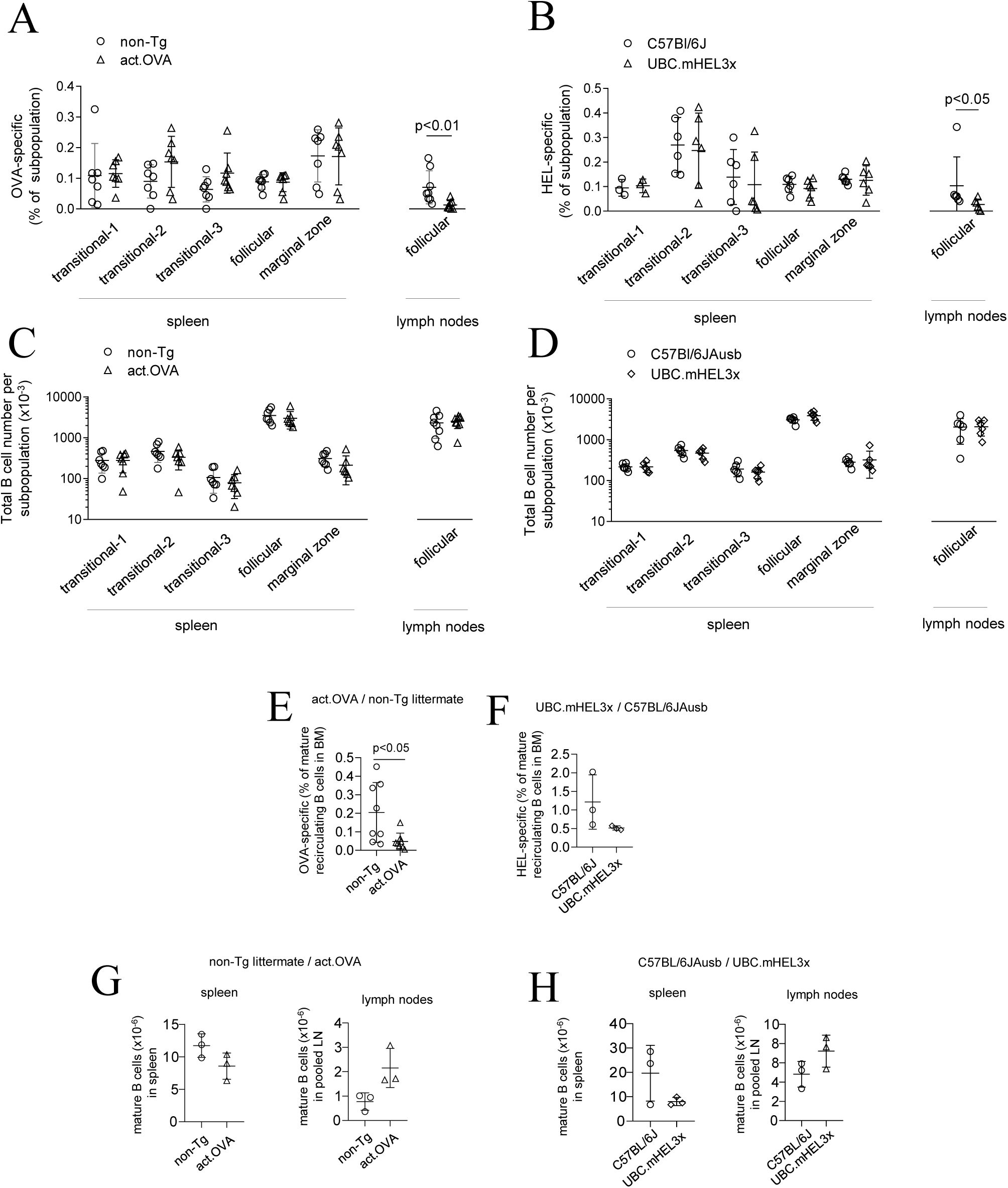
Characterisation of self-antigen specific B cell subpopulations in the spleen, lymph nodes and bone marrow and in aged mice. OVA and HEL tetramers were used to probe the spleen and lymph nodes of ‘young’ (A-F) or ‘aged’ act.OVA and UBC.mHEL3x mice (G,H). Antigen-specific B cells were enumerated among B cell compartments in each lymphoid organ using gating shown in Fig. 2A. A) Proportion of OVA-specific follicular, marginal zone, and transitional B cells among spleen and follicular B cell in lymph node of 10-14 weeks of age act.OVA mice and non-Tg littermate controls. B) Proportion of HEL-specific follicular, marginal zone, and transitional B cells among spleen and follicular B cell in lymph node of UBC.mHEL3x mice and C57BL/6JAusb controls C) Total number of follicular, marginal zone, and transitional B cells among spleen and follicular B cell in lymph node of act.OVA mice and non-Tg littermate controls. D) Total number of follicular, marginal zone, and transitional B cells among spleen and follicular B cell in lymph node of 10 weeks of age UBC.mHEL3x mice and C57BL/6JAusb controls. E,F) OVA- or HEL-specific B cells as a proportion of total recirculating mature B cells in BM. G) Total number of mature B cells in spleen and lymph nodes of aged (∼240 days) act.OVA mice and non-Tg controls. H) Total number of mature B cells in spleen and lymph nodes of aged (∼270 days) UBC.mHEL3x and controls. Data are values for individual mice pooled from 2 or more experiments. t-test with multiple testing correction and Mann-Whitney test (A,B) or t-test (E,F,G,H).

**Supplementary Figure 3.**
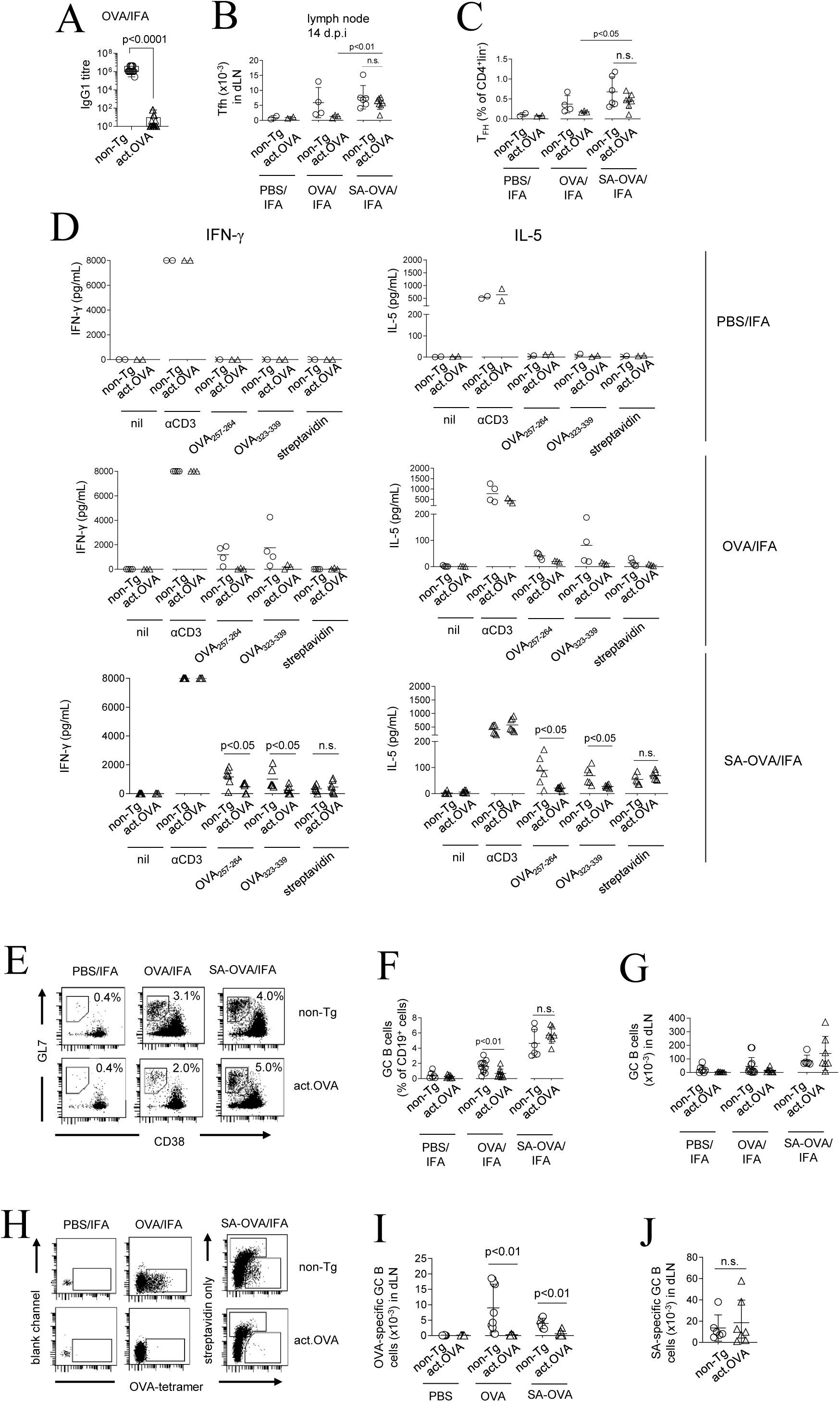
B-cell intrinsic mechanisms restrain self-reactive B cell function. Act.OVA mice and non-Tg littermates were immunized s.c. with PBS/IFA (sham), OVA/IFA or SA-OVA/IFA and analysed 14 days later (flow cytometry) or 14 days later (antibody production). A) OVA-specific IgG1 titres B) Number of Tfh in draining (pooled inguinal) lymph nodes 14 days after immunization. C) Frequency of Tfh in draining (pooled inguinal) lymph nodes 14 days after immunization. D) Concentration of IFN-γ or IL-5 in supernatant following in vitro restimulation (3 days) with antiCD3 (positive control), OVA_257-264_, OVA_323-339_, or streptavidin (SA), or left unstimulated (nil). E) Gating strategy and representative data for GC B cell analyses. F) Frequency of GC B cells in draining (pooled inguinal) lymph nodes 14 days after immunization. G) Total number of SA-specific GC B cell number in draining (pooled inguinal) lymph node 14 days after immunization. H) Gating strategy and representative data for antigen-specific germinal centre (GC) B cells. To separate SA-specific and OVA-specific GC B cells, inguinal lymph nodes were first stained with streptavidin-APC (to detect SA-specific B cells and to block SA-specific binding of subsequent tetramers) and then stained with OVA-PE tetramers. OVA and streptavidin-specific B cell gates were applied to GC B cell gates shown in D. Antigen-specific B cells were enumerated as the number of tetramer+ve cells_(unblocked – blocked)_ as described (Eur J Immunol 48: 1251-1254). I) Number of OVA-specific GC B cells in draining (pooled inguinal) lymph nodes 14 days after immunisation. J) Number of SA-specific GC B cells in draining (pooled inguinal) lymph nodes 14 days after immunisation. Figures show representative plots (D,G) or values for individual mice (mean±SD) pooled from 2 experiments (B,C) or 4 experiments (E,F,H,I). t-test (B,C,F) or Mann-Whitney test (I,J)

